# LazySlide: accessible and interoperable whole slide image analysis

**DOI:** 10.1101/2025.05.28.656548

**Authors:** Yimin Zheng, Ernesto Abila, Eva Chrenková, Juliane Winkler, André F. Rendeiro

## Abstract

Histopathological data are foundational in both biological research and clinical diagnostics but remain siloed from modern multimodal and single-cell frameworks. We introduce LazySlide, an open-source Python package built on the scverse ecosystem for efficient whole-slide image (WSI) analysis and multimodal integration. By leveraging vision-language foundation models and adhering to scverse data standards, LazySlide bridges histopathology with omics workflows. It supports tissue and cell segmentation, feature extraction, cross-modal querying, and zero-shot classification, with minimal setup. Its modular design empowers both novice and expert users, lowering the barrier to advanced histopathology analysis and accelerating AI-driven discovery in tissue biology and pathology.

## Main text

Histopathology is a cornerstone of biomedical research and clinical practice, offering high-resolution insights into the spatial organization of tissues, cellular morphology, and pathological alterations. Advances in digital pathology have enabled the collection of large repositories of whole-slide images (WSIs), creating unprecedented opportunities to study the tissue basis of human health and disease. However, computational analysis of WSIs remains challenging. Existing toolkits for WSI analysis such as QuPath^1^, CLAM^2^, PathML^3^, Slideflow^4^, and TIAToolbox^5^, offer partial solutions but are often limited by fragmented data structures, platform-specific constraints, and high technical barriers to entry. These limitations hinder integration with the multimodal and single-cell workflows that are increasingly standard in modern biology.

To address these limitations, we developed LazySlide, a general-purpose Python framework for WSI analysis, fully interoperable with the scverse ecosystem^6^ widely used in biomedical research. LazySlide streamlines WSI preprocessing, segmentation, feature extraction, and multimodal integration using state-of-the-art deep learning models – effectively bridging histopathology with omics workflows, data structures, and operations.

At its core, LazySlide introduces a custom data structure, WSIData (**Figure 1**), built upon scverse’s SpatialData^7^ but optimized for the diverse formats and scale of WSIs. While SpatialData supports only OME-TIFF pyramids, most WSIs are stored in proprietary formats requiring specialized readers. Existing extensions such as SOPA^8^ adapt SpatialData to WSIs but incur substantial disk overhead due to serialization (5-10x increase). In contrast, WSIData enables efficient, direct access to standard WSI formats without duplication, while maintaining compatibility with scverse tools and deep learning pipelines. The framework follows familiar API conventions from AnnData^9^, Scanpy^10^, and Squidpy^11^, reducing the barrier for computational pathologists and genomics researchers alike.

**Figure 1:**
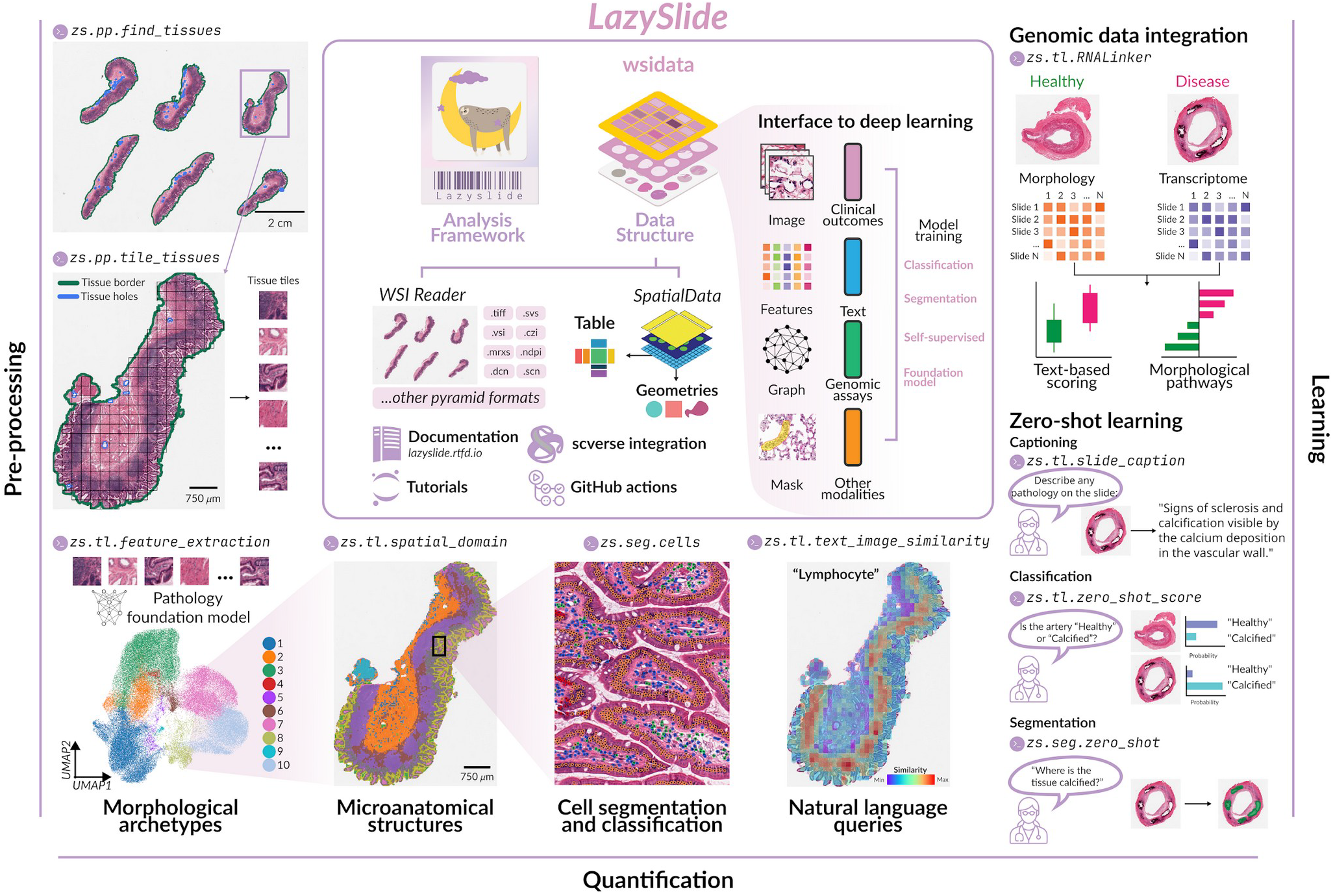
Overview of the LazySlide framework for scverse-compatible whole slide image analysis. Illustration of the capabilities of LazySlide enabled by its interoperable data structure WSIData: pre-processing of WSI data, quantification of cellular, morphological and microanatomical features of tissue, and learning by integration with other data types, interfacing with deep learning frameworks and advanced, multimodal capabilities.

LazySlide includes a full suite of tools for WSI analysis, from preprocessing, quantification, visualization, and multiomics integration, to advanced capabilities that support insightful interpretation and interactive exploration, leveraging zero-shot approaches and genomic data integration (**Figure 1**). Specifically, slides are segmented into tissue regions, tiled into memory-efficient patches, and embedded via vision foundation models into high-dimensional feature spaces. These embeddings support downstream analyses such as unsupervised clustering, which reveals spatial tissue architectures, for example in the human small intestine: mucosa, submucosa, muscularis, and lymphoid tissue (**Figure 1**). Built-in cell segmentation and classification tools enable quantification of cellular composition and morphology.

Beyond image-based analysis, LazySlide leverages vision-language models to support natural language queries, allowing users to localize image regions most similar to prompts like ‘lymphocyte’ for example. These features also enable multimodal integration with transcriptomics and other data types, supporting disease scoring and mechanistic interpretation. LazySlide further provides zero-shot capabilities for slide captioning, relevance scoring, and text-guided segmentation, as well as efficient, declarative visualization of WSIs and annotations, including segmented regions, cells, and extracted features.

LazySlide supports a diverse range of applications and facilitates rigorous benchmarking against existing methodologies, thereby showcasing its robust capabilities and broad utility. To showcase its capabilities in combining whole slide image (WSI) data with other modalities, we demonstrate three representative applications: i) zero-shot vision-language querying, ii) multimodal integration of histology and transcriptomics, and iii) zero-shot organ classification – each illustrating cutting-edge use cases for computational pathology and omics data integration.

First, we assembled a dataset of human artery slides from the GTEx project^12^, including healthy (n = 24) and calcified (n = 21) tissues with matched RNA-seq profiles (**Figure 2a**). With only five lines of code, users can compute text-to-image similarity maps across each slide. Terms related to “calcification” show higher enrichment in calcified samples, whereas anatomical terms predominate in healthy tissues (**Figure 2b-c**). A differential analysis of image-text features highlights terms such as gap junction, vascular niche, and apoptosis as significantly enriched in calcified arteries, consistent with observed morphological changes. Building on this, a slide-level “calcification score” is computed using top-k pooling over similarity maps with the term calcification (**Figure 2d**), yielding significantly elevated scores in calcified tissues.

**Figure 2:**
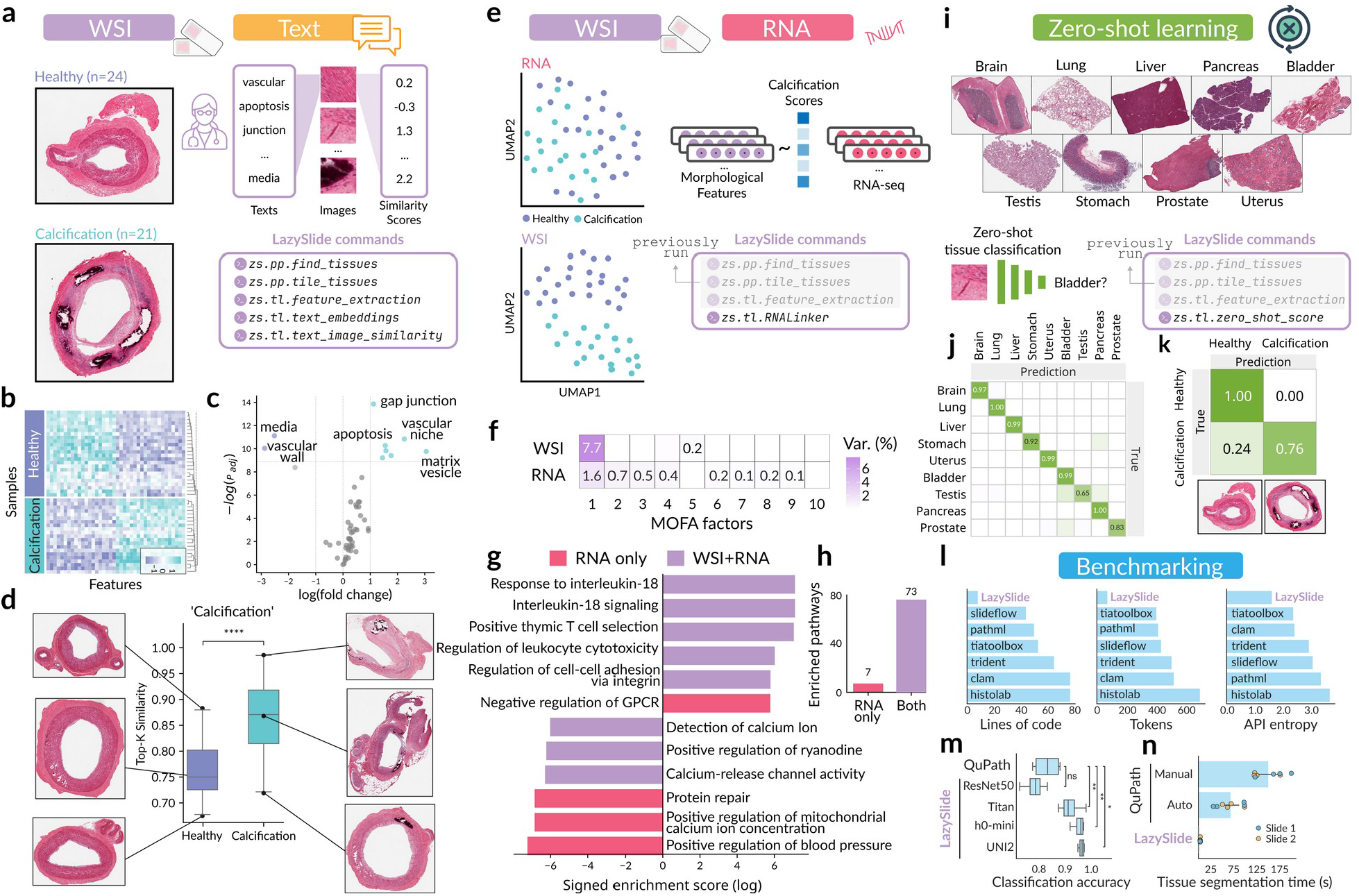
Diverse applications and performance benchmarks of LazySlide. **a)** Example whole slide images (WSIs) of healthy and calcified arteries, illustrating morphological differences which will be quantified via scoring of user-provided text terms from vision-text models. The full workflow can be run with 5 Lazyslide function calls. **b)** Heatmap with subset of text terms scored in each WSIs, illustrating a strong pattern associated with calcification. **c)** Discovery of differential associated text features between calcified and healthy arteries, highlighting key terms related to vascular health and disease. **d)** Quantitative scoring of arterial calcification for each WSI via text integration, with corresponding representative WSI tiles. p = 1e^-5^ (Mann-Whitney U test). **e)** UMAP visualization of RNA-seq data (top) or WSI data (bottom) from paired samples, showing clusters related to healthy and calcified states. Morphological features and RNA-seq can be linked for joint analysis. **f)** Variance explained by multi-omics factor analysis (MOFA) of RNA and WSI, indicating shared and unique biological variance. **g)** Enriched pathways identified through integrated WSI and RNA-seq analysis versus RNA-only analysis, demonstrating novel biological insights recovered by multimodal integration. **h)** Number of significantly enriched pathways from WSI+RNA integration versus RNA-only analysis, emphasizing the benefit of multimodal data. **i)** Representative examples from various tissue types used for zero-shot classification. **j)** Confusion matrix of zero-shot tissue classification predictions across different organs. **k)** Zero-shot prediction probabilities for calcification or healthy tissue, showcasing the ability to classify unseen pathological states. **l)** Comparison between LazySlide other frameworks, demonstrating LazySlide’s reduced code complexity. **m)** Benchmarking of various features of LazySlide against QuPath in classification accuracy. **n)** Benchmarking against QuPath in time to segment tissue from background, both in automated and manual workflows.

To explore joint imaging-transcriptomic patterns, we use the calcification score to anchor a multimodal integration of WSI-derived and RNA-seq embeddings (**Figure 2e**). This analysis, which is executable in a single line of code, shows that image features separate healthy and calcified groups more distinctly in UMAP space than RNA-seq alone. This pattern is also captured via multi-omics factor analysis (MOFA) integration using Muon, the scverse multi-omics framework^13^(**Figure 2f**). We then compared gene-level relevance to calcification by ranking genes from RNA-only differential expression or joint WSI+RNA integration. Pathway enrichment on the top 300 up- and down-regulated genes from each method revealed that only the WSI and RNA integration identifies key calcification-related pathways, including IL-18 signaling (**Figure 2g**), and yields a greater number of significant pathway associations overall (**Figure 2h**).

Finally, we demonstrate the zero-shot classification capabilities of LazySlide. Using vision-language models, WSIs from nine distinct human organs were queried against their organ names (**Figure 2i**). With a single line of code, LazySlide correctly identified the majority of organ sources (**Figure 2j**). Applying the same approach to the artery calcification dataset also yields high classification performance (**Figure 2k**), which illustrates the ability to extract valuable insights without tissue- or context-specific training.

To assess usability and performance, we benchmarked LazySlide against established tools across a standard preprocessing pipeline: tissue segmentation, tiling, tile dataset preparation for PyTorch, and feature extraction. LazySlide completes this workflow with fewer lines of code, lower token count, and a simpler API, facilitating rapid development and code maintenance (**Figure 2l**). We also compared LazySlide to QuPath^1^ in a classification setting of semantically labeled microanatomical domains of murine breast cancer lung metastasis using extracted features by different vision models. Excluding ResNet50 which is not trained on tissue images, LazySlide consistently outperforms QuPath-derived features (**Figure 2m**). Moreover, tissue segmentation is markedly faster in LazySlide compared with QuPath using either automated or manual workflows (**Figure 2n**).

Altogether, LazySlide represents a significant advancement in AI-enabled histopathology and tissue biology, bridging computational pathology with multimodal omics through a modular, user-friendly, and open-source framework. By supporting a wide range of foundational models and ensuring seamless interoperability with the scverse ecosystem, LazySlide enables integrative, scalable, and interpretable analysis of whole slide images. It empowers both computational pathologists and genomics researchers to uncover new insights into tissue biology and disease, accelerating the development of data-driven, clinically meaningful models.

## Online methods

### Data structure

The WSIData object was designed as an inheritance of SpatialData^7^, allowing it to behave exactly like SpatialData. This design enables users familiar with scverse and SpatialData to readily use WSIData across scverse packages such as Lazyslide. The SpatialData component of WSIData is a data container, storing all analysis metadata and results generated within LazySlide. An additional layer and corresponding APIs were implemented to specifically interact with whole slide images. WSIData was built to support multiple WSI backends, including OpenSlide^14^, tiffslide, bioformats^15^, and cuCIM, ensuring compatibility with a broad range of image formats. An accessor interface, similar to those found in pandas, xarray, and SpatialData, was also implemented to provide a variety of features. For instance, users can easily retrieve useful information, such as the number of tissue pieces found in each slide, the number of tiles in a tesselated image, and an AnnData object ^9^ with morphological features through a ‘*fetch*’ accessor. The ‘*iter*’ accessor allows users to iterate through each tissue piece, tile, or their features. The ‘*dataset’* accessor provides an interface to create various PyTorch^16^ datasets that could be directly used for training or running deep learning models.

### Tissue segmentation

Tissue segmentation was performed using a multi-step image processing pipeline, which was implemented with OpenCV. The algorithm initiates processing of WSIs at an automatically determined optimal resolution level. For artifact filtering, specialized thresholding techniques were applied to identify and exclude non-tissue regions based on color properties. Alternatively, images can be either converted to grayscale or transformed to HSV color space, with the saturation channel extracted to enhance tissue visibility. A median blur filter was first applied to reduce noise, followed by binary thresholding, either using Otsu’s method or a fixed threshold value. The resulting binary mask undergoes morphological closing operations to fill small gaps within tissue regions while preserving overall tissue structure. This process effectively separated tissue regions from background and artifacts, enabling subsequent region identification and analysis. Tissue masks can be further refined using higher-resolution image data when specified, and regions below a configurable minimum area threshold are excluded. As an alternative, the GrandQC^17^ pre-trained tissue segmentation model was also integrated into LazySlide to perform deep learning-based tissue segmentation.

### Tissue tiling

Tissue regions were tiled using an automated approach that generated a grid of tiles within identified tissue masks/contours. The algorithm allows users to request arbitrary tile sizes no lower than the raw image resolution and accepted user-specified tile dimensions along with customizable stride or overlap between adjacent tiles. For each tissue region, tiles were initially generated within its bounding box, then filtered to retain only those intersecting with the tissue contour. To eliminate tiles containing excessive background, a configurable background fraction threshold is applied, discarding tiles where the background exceeded this threshold. The tiling process accommodates multi-resolution analysis by allowing the user to specify a resolution in microns-per-pixel for consistent physical scale across slides scanned at variable magnifications.

### Quality control

Following tiling, quality control was implemented through a configurable scoring system that evaluated tiles based on metrics including focus quality, contrast, brightness, and tissue characteristics. This scoring facilitated subsequent filtering of low-quality tiles containing artifacts or out-of-focus regions, thereby ensuring the reliability of downstream analysis. All operations supported parallel processing for computational efficiency when handling large datasets. Additionally, GrandQC^17^ was incorporated in LazySlide to detect common artifacts such as bubbles, tissue folds, and out-of-focus regions.

### Feature extraction and aggregation

The feature extraction module supports a variety of pre-trained vision models through Torch Image Models (timm, https://github.com/huggingface/pytorch-image-models). The tiled images are processed through selected models, including classic vision architectures pre-trained on ImageNet (e.g. ResNet, DenseNet, ViT) and pathology-specific foundation models (e.g. UNI/UNI2^18^, Virchow/Virchow2^19,20^, H-Optimus-0/1, GigaPath^21^), to extract high-dimensional feature representations. Feature extraction supports automatic device selection (CPU/GPU), mixed-precision inference, and configurable batch processing for computational efficiency. For each tissue tile, the deep learning model extracts feature vectors that capture complex visual patterns not easily quantifiable through traditional image analysis. Following extraction, LazySlide implements multiple feature aggregation strategies to generate slide-level or tissue-level representations from tile-level features. These include statistical aggregation methods (mean, median) and specialized neural encoders that incorporate spatial information (e.g. PRISM^22^, TITAN^23^).

### Natural language query of tissue images

Natural language query capabilities in LazySlide leverage multimodal foundation models to bridge vision and language domains, enabling content retrieval through text-based queries. The framework employs state-of-the-art pathology-specific models, such as PLIP^24^ and CONCH^25^, to facilitate semantic search within whole slide images. The query process follows a two-stage approach: first, text queries are transformed into dense vector representations using the text encoder component of the chosen model. Second, these text embeddings are compared with pre-extracted image feature vectors using cosine similarity to identify regions most relevant to the query. Similarity scores are computed via the dot product between normalized text and image embeddings, with higher scores indicating stronger semantic correspondence. This powerful functionality enables users to search for specific histological patterns, cell types, or tissue structures using natural language descriptions, offering a significant advantage over traditional visual inspection methods.

### Spatial domain detection

LazySlide implements unsupervised spatial domain identification through a multi-stage, graph-based clustering approach. Initially, a neighborhood graph is constructed from the extracted tiles, where each tile represents a node and edges connect spatially adjacent tiles. Optionally, morphological features can be smoothed by incorporating neighboring information, thereby enhancing feature consistency. Following graph construction, LazySlide employs a dimensionality reduction pipeline, as introduced in Scanpy^10^, which first scales the feature data and then applies principal component analysis to capture dominant variation patterns. The resulting low-dimensional representation is then used to construct a weighted k-nearest neighbor graph. Finally, the Leiden community detection algorithm (implemented via igraph) identifies coherent tissue domains by partitioning this graph into communities, with the clustering resolution controlled by a user-adjustable parameter. This comprehensive approach effectively segments histologically distinct regions within tissue slides without requiring manual annotation, thereby enabling automated identification of intricate tissue architectural features.

### Whole slide cell segmentation and classification

LazySlide supports cell segmentation of whole slide images through Instanseg^26^ and joint cell segmentation with classification through Nulite^27^. The segmentation stages operate within a dedicated SegmentationRunner class, which efficiently handles batch processing across tiles, utilizing GPU acceleration when available, and performs the crucial merging of results from individual tiles. For each tiled image, cells are identified and their contours are transformed into a polygon representation. To merge results across tiles of the whole slide image, results are aggregated by addressing potential boundary artifacts, via a sophisticated polygon merging algorithm that leverages a spatial indexing tree (STR tree). All initially overlapping polygons are identified and then merged by preserving their class labels while calculating probability-weighted averages for cells spanning multiple tiles, thereby ensuring spatial continuity across the entire tissue section. Newly merged polygons are then pruned from the tree, and the process iterates until the tree contains no remaining branches, ensuring all overlapping polygons have been successfully merged into cohesive cell instances.

### Integration with RNA-seq data

LazySlide enables multimodal data integration through the RNALinker class, which links histopathological features with paired transcriptomic profiles. Morphological features extracted from whole slide images are first aggregated into slide-level representations that are matched to corresponding genomic samples. The integration proceeds in two steps: 1) samples are grouped and scored based on morphological features – either through differential analysis between defined sample groups or by using morphologically-derived annotations, such as text-based scores; 2) these morphology-based scores are statistically associated with gene expression data using correlation metrics (e.g., Pearson, Spearman, Kendall) and regression models (e.g., linear regression, lasso). This approach identifies genes whose expression is significantly associated with specific histological patterns. By linking morphology to transcriptomic signatures, this integration supports hypothesis generation about the molecular mechanisms driving tissue phenotypes and facilitates the discovery of biomarkers that bridge computer vision-derived features with biological interpretation.

### Zero-shot learning

LazySlide supports zero-shot learning through multimodal vision-language foundation models such as PRISM^22^ and TITAN^23^, supporting various biologically relevant questions without task-specific training. For zero-shot classification, slide-level feature embeddings are compared to arbitrary, user-defined text prompts, yielding probability scores that quantify the semantic alignment between tissue morphology and natural language descriptions. Alternatively, for descriptive analysis, LazySlide generates natural language summaries of histological content from slide embeddings and text prompts, enabling automated preliminary reporting and assisting pathologists with concise, context-aware interpretations. For zero-shot segmentation of tissue, LazySlide uses text-image similarity metrics to produce a binary mask based on a user-defined threshold. Detected object bounding boxes from this mask are then used to guide segmentation via the Segment Anything Model 2 (SAM2)^28^, enabling flexible and label-free identification of tissue regions of interest.

### Benchmarking

To assess the usability and interoperability of LazySlide within the scverse ecosystem^6^, we conducted a benchmark comparison against several widely used whole slide image (WSI) processing libraries: CLAM^29^, TRIDENT^30^, PathML^3^, Tiatoolbox^5^, Histolab^31^, and Slideflow^4^. Each library was evaluated using a standardized digital pathology workflow comprising tissue segmentation and tiling, feature extraction with ResNet50, construction of a PyTorch dataset for tile-level access (to assess ease of integration with deep learning), and generation of an AnnData object containing tile features and spatial metadata (to assess compatibility with scverse workflows). Benchmark scripts were implemented for each library, formatted using *ruff*, and executed in isolated Docker environments to ensure consistency and reproducibility. To quantify implementation complexity, we measured token counts, lines of code, and API entropy, providing a comparative view of conceptual and practical overhead.

For direct benchmarking against QuPath, we used four mouse lung tissue slides from a breast cancer patient-derived xenograft (PDX) model with metastasis. These slides were previously semantically annotated for three tissue classes: tumor, airways, and blood vessels. Feature extraction was performed using both QuPath^1^ and LazySlide (with ResNet50, h0-mini, UNI2, and TITAN backbones), and classification accuracy was used for performance comparison. Tissue segmentation was benchmarked by comparing QuPath (manual and automated methods) with LazySlide. Following a standardized protocol, four experienced volunteers used QuPath’s manual labeling and thresholding tools, and the time needed to perform tissue segmentation was recorded for two slides. The time performance of LazySlide was measured using the *find_tissues* function.

## Data availability

GTEx whole slide images and RNA expression data are available from its portal (*https://gtexportal.org*). Whole slide images of a breast cancer PDX models and their semantic segmentation are available at Zenodo (https://doi.org/10.5281/zenodo.15497223).

## Code availability

Lazyslide and WSIData are publicly available as GitHub repositories (https://github.com/RendeiroLab/LazySlide, https://github.com/RendeiroLab/WSIData), published on the Python package index (PyPI), deposited on Zenodo (https://zenodo.org/records/15083451, https://doi.org/10.5281/zenodo.15083425), and documented online (https://lazyslide.readthedocs.io, https://wsidata.readthedocs.io). A companion repository on benchmarking is also publicly available on Github (https://github.com/RendeiroLab/LazySlide-benchmark). We thank Elisabeth Weigert, Elisabeth Gurnhofer, Patrick Wagner, and Gerald Timelthaler for their technical support and pathologists Zsuzsanna Bagó-Horváth and Ulrike Heber for their feedback on tissue annotations.

## Acknowledgments

This research was funded by a grant from the Vienna Science and Technology Foundation (WWTF-LS23-067). A.F.R was supported by Angelini Ventures S.p.A. Rome, Italy. The Genotype-Tissue Expression (GTEx) Project was supported by the Common Fund of the Office of the Director of the National Institutes of Health, and by NCI, NHGRI, NHLBI, NIDA, NIMH, and NINDS. We thank all Rendeiro Lab members for their precious input and for testing LazySlide.

## Author contributions

Y. Z. and A.F.R conceptualized the study; Y.Z., led the development of with contributions from E. A.; E.C. provided annotations for WSIs; J.W. and A.F.R. supervised the research. All authors reviewed and approved the manuscript.

## Conflicts of interest

The authors declare no competing financial interests.

## Notes

https://github.com/rendeirolab/LazySlide

https://lazyslide.readthedocs.io/

https://github.com/rendeirolab/WSIData

https://wsidata.readthedocs.io/

### Competing Interest Statement

The authors have declared no competing interest.

